# Daily feeding rhythms may play a role in the genetic variability of feed efficiency in growing pigs

**DOI:** 10.64898/2026.04.17.719142

**Authors:** Hélène Gilbert, Aline Foury, Lawal Agboola, Guillaume Devailly, Florence Gondret, Marie-Pierre Moisan

## Abstract

Improving feed efficiency in pigs is essential for reducing production costs and environmental impacts. This study examines the influence of circadian feeding rhythms and genetic polymorphisms on feed efficiency variability using two pig lines divergently selected for Residual Feed Intake (RFI) over ten generations. Feeding behavior was monitored using automatic concentrate dispensers, recording 6,494,097 visits from 3,824 pigs to analyze meal frequency, duration, and diurnal patterns. LRFI pigs ate less frequently, with larger meals and longer durations, they exhibited two distinct feeding peaks: one around 8:00 AM and a higher one at 5:00 PM and they consumed more feed during the diurnal period and less at night. HRFI pigs showed a smoother, less rhythmic feeding behavior with increased nocturnal intake. The differences between the two RFI lines became more pronounced as the number of generations of selection increased, suggesting a genetic basis. Feeding behaviors, including intake during the two main diurnal peaks, were found to be heritable (heritability estimates: 0.30-0.40) and genetic correlations were observed between feed intake and RFI, especially for intake between the two peaks. Then, we investigated the evolution of allele frequencies of single nucleotide polymorphisms (SNPs) in DNA sequences surrounding 10 core clock genes (*ARNTL, CLOCK, CRY1, CRY2, NPAS2, NR1D1, PER1, PER2, PER3, RORA*) along generations of selection. SNPs with significant frequency changes were mapped to regulatory regions and transposable elements, especially in HRFI line, suggesting potential functional impacts on circadian regulation. These results underscore the role of feeding behavior and genetic variation in feed efficiency, offering insights for breeding programs aimed at improving metabolic efficiency and sustainability in pig production.

## Introduction

Circadian rhythms are central to physiological regulations in bacteria, plants, and animals including humans, influencing metabolism, behaviour and biological functions (Manoogian et al., 2022; Panda, 2016; Potter et al., 2016). Alignment of feed intake with day/night cycles is critical for the regulation of energy metabolism and body composition as shown by numerous studies in rodent models and humans (Boege et al., 2021). In pigs also, the disruption of circadian rhythms affected energy metabolism, with pigs fed at night instead of during the day having increased retention of energy in the form of fat (van Erp et al., 2020). Circadian rhythms are regulated by biological clocks, which are molecular systems remarkably conserved throughout evolution. The main clock is located in the suprachiasmatic nucleus of the hypothalamus in the brain. It is synchronized by natural light and regulates the biological clocks of all other tissues, according to autonomous endogenous molecular oscillations, which are determined by a limited set of genes called core clock genes (Takahashi et al., 2008).

The objective of this study was to evaluate whether variations in the circadian rhythms, identified through variations in the feeding patterns of thousands of pigs during night and day, were associated with variations in feed efficiency. Feed efficiency is the ratio of average daily weight gain to average daily feed consumption over a given period. An indicator of feed efficiency is residual feed intake (RFI), which is defined as the difference between actual feed intake and expected feed intake for maintenance (predicted from metabolic body weight) and production performance (predicted from weight gain and composition) (Gilbert et al., 2017). Previous studies in divergent RFI lines have revealed that difference in feed efficiency was underlined by variations in transcripts of many genes participating in several functional pathways (Gondret et al., 2017), so that feed efficiency is a complex polygenic trait. Phenotypically, the low RFI pigs (LRFI), i.e. the most efficient, were leaner and energy metabolism was also different as compared to high RFI pigs (HRFI): glycogen utilization was up-regulated whereas fatty acid oxidation was down-regulated in muscle of the LRFI pigs as compared to the HRFI pigs. When analyzed thanks to video recordings of the behavioral activity during growth-finishing period, it has been shown that LRFI pigs also spent less time eating, with shorter daily eating time, lower number of visits and increased feeding rate during the day than the HRFI pigs, (Meunier-Salaün et al, 2014). To date, eating behaviour among pigs can be tracked using farming technologies such as automatic concentrate dispensers, which reveals diverse behaviors (daily meal eaters, nibblers, day-night eaters) and several inconsistencies from day to day (Bus et al., 2023). Assuming that part of these differences could result from changes in circadian rhythms, the present study aimed to investigate changes in eating patterns throughout day and night in a large number of growing pigs from these two divergent lines for RFI and along ten generations of selection. Then, the variance components of the eating rhythm characteristics were estimated to quantify the genetic bases of their variability, and genetic correlations with feed efficiency traits were assessed. The changes of allele frequency of loci of the core clock genes along selection were tested in the two RFI lines, and functional effects of the identified SNPs were investigated.

## Material and methods

### Ethic statement

The care and use of pigs were performed following the guidelines edited by the French Ministries of High Education and Research, and of Agriculture and Fisheries (http://ethique.ipbs.fr/sdv/charteexpeanimale.pdf). All pigs were reared in compliance with national regulations and according to procedures approved by the French Veterinary Services.

### Animals

Data from 3,991 pigs from generations G0 (first generation of selection) to G9 (tenth generation of selection) of two lines of pure Large White pigs divergently selected for RFI were used; ie all data available from these experimental lines. The selection began in 1999 in the experimental breeding INRAE unit GenESI (Surgères, France, https://doi.org/10.57745/LN1IZE). Principles of selection were described by Gilbert et al., 2007. Briefly, daily feed intake (DFI; kg/d) was measured on group-housed boars fed ad libitum from 35 to 95 kg body weight (BW). Two traits were recorded and used to calculate the predicted feed intake: average daily gain (ADG, kg/d) from 35 to 95 kg BW and backfat thickness (BFT, mm) at 95 kg BW assessed by an ultrasonic device and averaged from 6 different measures on the back. A phenotypic RFI selection index was then computed as a linear combination of those traits as follows: RFI = DFI -(1.06 × ADG) -(37 × BFT). The weights assigned to ADG and BFT in the selection index were chosen to keep ADG and BFT constant at the phenotypic level between the two RFI lines. Since RFI is the difference between measured and predicted feed intake for pig production and maintenance requirements, a negative RFI corresponds to pigs that ingest less feed than predicted. Therefore, low RFI pigs (LRFI) were more efficient than high RFI pigs (HRFI line). At each generation, 96 male pigs per line were measured for RFI, and the best 6 pigs in each line were retained to produce the next generation. There was no selection pressure on the female pathway, with about 40 females kept per line and one gilt replacing its dam.

The pigs were kept in pens of 12 pigs, by sex (female, male or castrated male) and line (LRFI, HRFI), and by contemporary batches (183 batches of animals in total). Each contemporary group comprised pigs born during the same week from both lines. Body weight at birth was 1,588 ± 350 grams and weaning BW was 8,893 ± 2,105 grams. Performance measurements were carried out between 10 weeks of age and commercial slaughter weight (115 to 120 kg BW). One electronic feeder (ACEMO 64 SEF, Acema, Pontivy, France) was present in each pen of group-housed pigs. The pigs had free access to water and a commercial pellet feed formulated for growing-finishing pigs, containing an average of 10 MJ of net energy (NE) and 160 g of protein per kg during the test. The feed intake was available daily thanks to the electronic feeder. The ADG and average DFI during the test period were calculated, as well as average metabolic weight (AMW, Noblet et al., 1999). The realized residual average daily feed intake was then estimated as the residual of the multiple linear regression of the DFI on ADG, AMW and BFT. The RFI computed for the different sex (male, castrated male or female) were combined as described in Aliakbari et al. (2020).

### Data cleaning and feeding rate

Feeding activities were recorded for each pig during the 3 months of test, starting at 10 weeks of age until 115 to 120 kg BW. When a pig entered into the electronic feeder, the door closed behind it, allowing one individual animal to eat at each time. An ear tag with an RFID transponder allowed identification of the pig in the feeder. The quantity of feed consumed, and the time when the feeding started and ended were recorded. The raw data were the 7,066,642 visits to the feeders recorded for 3,991 animals. Prior to analysis, visits without any feed ingestion (< 0 g) or with negative feeding durations were considered aberrant and removed from the dataset. The few pigs with less than 65 days with feeding records were also removed from the analysis. This resulted in 6,410,928 visits for a total of 3,824 pigs.

Afterwards, two complementary variables including visit duration, time spent per visit (TSV) and time interval between visits (IBV) were computed for each visit. The visit duration is the difference between the time pig left the feeder and the time it entered. The interval between two visits was calculated as the difference between a visit’s start time and the previous visit’s end time. Then, visits that had a negative interval with the previous visit were set to missing. Lastly, the intervals between visits that lasted for more than 24 hours and less than 20 seconds were also set to missing. These unusual records might be due to potential issues with the electronic feeders. The distributions of derived feeding patterns traits (quantity fed per visit, visit duration, intervals between visits, feeding rate, and the number of visits per day) were then visually checked. They were all far from normality and contained extreme values. Then, outliers were removed by eliminating all values of observations that were out the range of ±4 standard deviations of their mean in all traits after applying a log transformation to get closer to normality.

To describe the distribution of the feeding rhythm over a 24-hour period, the average quantity ingested per hour was then calculated for each pig, i.e. 24 traits. In addition, the average quantities of feed ingested during the day (06:00 to 21:00) and during the night (21:00 to 06:00) were calculated. Finally, because feeding rhythm is reputed to be mainly distributed in two peaks during the day, one in the morning and one in the afternoon (Nielsen et al., 1995; Renaudeau, 2009) the average feed intakes during the first peak (06:00-10:00) and during the second peak (15:00-21:00) were also calculated for each pig, as well as the average intake between the peaks (10:00-15:00).

### Molecular data

All breeding stocks from the low and high RFI lines were genotyped with chips containing around 60,000 markers (Delpuech et al., 2021). In addition, the 20 G0 animals that contributed most to the selection generations (the 2 x 6 sires plus 8 dams) were sequenced at an average depth of 25X (Illumina short-read WGS). These individuals, along with 12 others major G0 dam contributors, were genotyped using the 650K chip (Affymetrix Axiom Porcine Array). The sequences were analyzed using the nf-core/sarek workflow to align the sequences with the reference porcine genome (Sscrofa11.1) and list all SNP variants. A total of 20,494,824 SNPs was identified and filtered. To obtain genotypes for all individuals at all these positions, sequential imputation was carried out (imputation of 650K markers from 60K genotypes, then imputation of 15,847,620 SNPs from the 650K imputed genotypes) using FImpute3 software (Sargolzaei et al., 2014). The correlation between true genotypes and imputed genotypes was used to evaluate imputation quality, with a score of 0.94 on the G0 individuals.

In the coding sequence of 10 core clock genes (*ARNTL* – also known as *BMAL1* –, *CLOCK, CRY1, CRY2, NPAS2, NR1D1, PER1, PER2, PER3* – ENSSSCG00000028572 –, *RORA*), 5,924 SNPs were identified as polymorphic in the two lines. When the region has been extended to 10^6^ bp on either side of the coding sequence, 145,448 SNPs were found in our dataset. The genotypes of the SNPs were extracted for all breeders in both lines. The SNPs were identified by the chromosome number on which they are located and by their position on the chromosome.

### Statistical analyses

#### Differences in feeding patterns between lines

To estimate the effects of selection on feeding traits, linear models were successively applied to all calculated traits to evaluate the effect of the line at the different time points considered (per hour or per period): **Y** = µ + **Xβ** + **e**, where **Y** is the vector of feeding traits, **β** comprises the fixed effects of the contemporary group within generation (n=183), sex (n=3), line (n=2), pen (n=16) and time (n=24 for the hours; n=2 for the other periods) were used. The interaction line x generation was fitted, as well as line x generation x time. In some models, DFI was added as a covariate to adjust traits for differences in feed intake between animals. **X** was the incidence matrix for the fixed effects, and **e** was the random residual. When hourly data were analyzed, the additional random effect of the repeated animal across hours was included in the model. The R software, R version 4.1.2 (R Core Team, 2021) was used, with packages “car” (Fox and Weisberg, 2019) for the linear models, and “lmerTest” (Kuznetsova et al., 2017) for the linear mixed models, were used for these analyses. Given the large sample sizes and trait variances, feed intake differences below 50g/h (3% difference between lines) could be detected with a 5% type I error.

#### Variance components estimations

Univariate linear mixed models with the same fixed and random effects as in the previous section, plus a random additive genetic effect, with a normal distribution of mean zero and variance-covariance 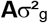, where **A** is the pedigree kinship matrix and 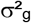 the genetic variance, were applied to all traits to estimate their variance components. The pedigree comprised all the generations of selection of the lines, plus 5 generations of ancestors, for a total of 6,834 individuals. Bivariate models were then used to estimate their genetic correlation with RFI and DFI. These analyses were carried out using ASRemL 3.0 software (Gilmour, 2019).

#### Evolution of allelic frequencies with selection

For each of the studied SNPs, the allelic frequencies in each line and generation were calculated using PLINK software (www.cog-genomics.org/plink/1.9/, Chang et al., 2015). Within-line, a regression of the number of generations of selection on the allele frequency of the minor allele of the population was then applied to each SNP, to detect significant changes in allele frequencies during selection (R software, “lm” function). To take into account of the fact that the genome contains several million SNPs and that multiple tests are applied, a test statistic threshold was calculated by adjusting for the corresponding number of independent tests at the genome level (package R SimpleM, Gao, 2011), leading to a genome-wide threshold of 7.22 for the -log_10_ of the p-value at 5% of each regression test. Given the frequency variations from one generation to the next and the genome-wide threshold applied, this approach allowed detecting linear frequency changes larger than 0.041 per generation, ie only large frequency changes over the ten tested generations were detected. The average chromosome-wide threshold was 5.95, rounded to 6 in the following analyses. Any signal significant at the chromosome-wide level was considered as suggestive.

### Predicting SNPs functional effects

#### Predicted effects of the SNPs

To predict the effects of SNPs identified as at least significant at a chromosome-wide threshold (-log_10_(P-value) > 6), the Ensembl Variant Effect Predictor was used (VEP, McLaren et al., 2016). Analysis was run with the following parameters: Pig reference (*Sus_scrofa*) as the species; a Variant Call Format file (VCF) for the 145,448 SNPs as the input data and Ensembl/GENCODE and RefSeq transcripts as the transcript database. We considered as SNPs of interest SNPs with one of the following effects: 3_prime_UTR_variant (a UTR variant of the 3’ UTR), 5_prime_UTR_variant (a UTR variant of the 5’ UTR), downstream_gene_variant (a sequence variant located 3’ of a gene), missense_variant (a sequence variant, that changes one or more bases, resulting in a different amino acid sequence but where the length is preserved), non_coding_transcript_exon_variant (a sequence variant that changes non-coding exon sequence in a non-coding transcript), non_coding_transcript_variant (a transcript variant of a non-coding RNA gene), start_lost (a codon variant that changes at least one base of the canonical start codon), stop_gained (a sequence variant whereby at least one base of a codon is changed, resulting in a premature stop codon, leading to a shortened transcript), stop_lost (a sequence variant where at least one base of the terminator codon (stop) is changed, resulting in an elongated transcript), stop_retained_variant (a sequence variant where at least one base in the terminator codon is changed, but the terminator remains), upstream_gene_variant (a sequence variant located 5’ of a gene).

#### Identifying SNPs in key regulatory patterns

For miRNA binding regions, the miRDB target prediction database was used to identify miRNA binding regions in clock genes 3’UTR region (https://mirdb.org/, Chen and Wang, 2020). Cry2 was the only clock gene with a SNP in the 3’UTR region. As pig was not one of the species available in the database, we used the human data since pig and human *CRY2* coding sequence display 74% homology.

For transposable Elements DNA sequences, the Dfam database was used to identify if SNPs were located in Transposable Elements DNA sequences (TE, https://dfam.org/, Storer et al., 2021). The Sequence Search tool was used with Sus scrofa as the source organism.

#### SNPs conservation level and localization in regulatory regions

The 1,886 SNPs significant (-log_10_(P-value) > 6) for the evolution of allele frequencies and with outcome in the VEP analysis were selected. Co-localization of statistically significant SNPs with regions of high conservation or with regions of putative regulatory regions as defined by Ensembl v111 (Harrison et al., 2024) were examined with three lists of conserved regions, as defined by comparative genomics approach on a set of 16 pig breeds, 63 amniotes, or 91 mammals. Regulatory elements were defined by Ensembl using GeneSwitch ChIP-seq dat of CTCF and Histone post-translational modification and ATAC-seq data. Co-localisations were performed using bedtools (Quinlan and Hall, 2010) and visualized using UpSetR (Conway et al., 2017). Transcription factor motifs were taken from JASPAR 2022 (Castro-Mondragon et al., 2022), using the vertebrate set. For each motif of size N, we tested a sequence of length N+2 centered on each SNP using the searchSeq function of the TFBS Tools R package (Tan and Lenhard, 2016) using a minimum hit score of 90%.

## Results

### Averaged daily feeding pattern traits

When averaged across days, sexes and generations of selection, all line differences at the day and visit levels were different (P < 0.0001, Table 1). Daily feed intake and feeding time were lower for LRFI pigs than for the HRFI pigs, associated with a lower number of visits to the feeder per day; however, the LRFI pigs ate more per visit and for a longer time, resulting in a higher feeding rate.

**Table 1.**
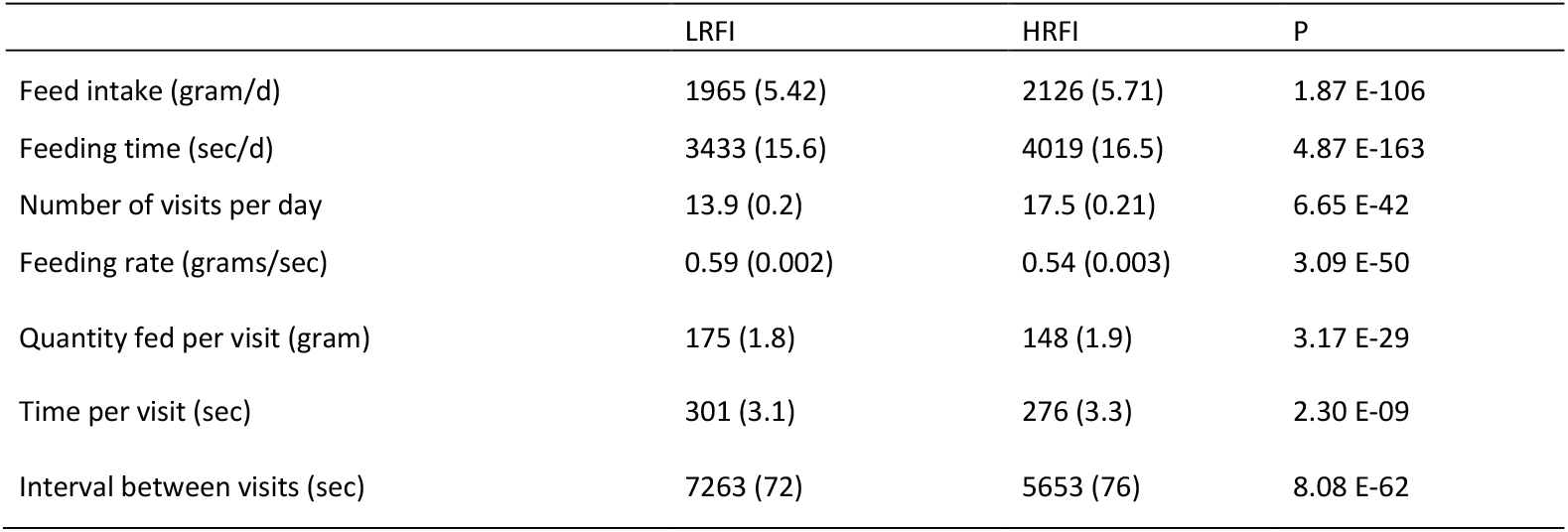
Least Square Means and the standard error of effect of line on the feeding pattern traits per day and per visit.

### Average feed intake per hour

The distribution of average feed intake per hour during the day for each line is shown in Figure 1, for all generations (panel A) and for the last generation (tenth) only (panel B). The general pattern for the feed intake distribution included two peaks of feed ingestion during the day, one around 08:00 and one around 18:00. These peaks were sharper in LRFI pigs than in HRFI pigs, with higher feed intake during the peaks and lower feed intake between peaks for the LRFI line. During the night (21:00 to 06:00), there was 22% of the total feed ingested in the two lines. When all generations were averaged, the LRFI pigs consumed less feed than the HRFI pigs between 22:00 and 06:00 (P < 0.001), and from 09:00 to 16:00 (P < 0.05). The amount of ingested feed was similar in the two lines when analyzed in the middle of the first peak. However, the LRFI pigs ingested more than the HRFI pigs between 18:00 to 21:00 (P < 0.05). Focusing on the last generation of selection, these differences were even more pronounced; in addition, in this last generation of selection, the LRFI pigs also ingested more between 06:00 and 08:00 (P < 0.05). When adjusted for the total individual feed intake of the pig (panels C and D), the differences during the peaks increased, showing a higher relative intake for the LRFI pigs during those, whereas the night differences were reduced when considered relatively to the total intake of the animals.

**Figure 1.**
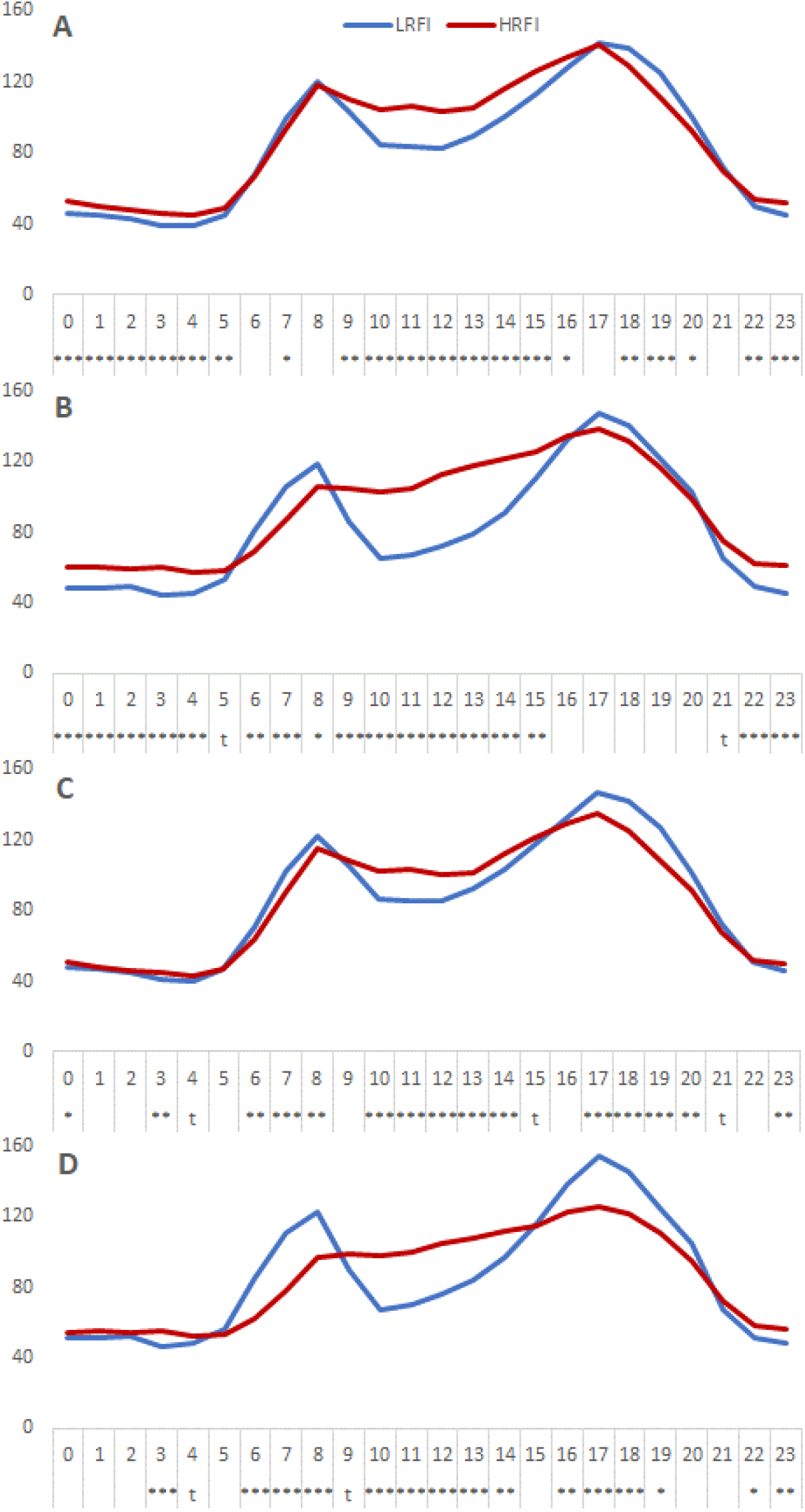
Average feed intake in grams per hour and per pig during the day for each line for all generations combined (A), for the last generation (B), for all generations combined adjusted for individual daily feed intake (C) and for the last generation adjusted for individual daily feed intake (D). Significance levels of line differences for each hour are shown below the x-axis. t (tendency) = P <0.10; * = P < 0.05; ** = P < 0.01; *** = P < 0.001.

### Distribution of average diurnal and nocturnal feed intake along selection

Table 2 reports least-squares means for each line x generation combination for diurnal and nocturnal intakes, unadjusted or adjusted for daily feed intake (DFI). The unadjusted models show significant differences in feed intake between pigs of the LRFI line and pigs of the HRFI line, from generation G3 (diurnal) and G4 (nocturnal) onwards, with consistently lower feed intakes in the LRFI line. However, after adjustment for DFI, there were no differences between the two lines until G6. From generation G7 onwards, there were higher feed intakes in the LRFI line during the day, and reduced feed intakes at night (P < 0.01) compared with the HRFI line.

**Table 2.** Average daytime and overnight intakes for each line and generation (G0 to G9) in g/period. Least squares means of the line x generation effect in a linear model with (adjusted) or without (unadjusted) average daily intake as covariate, standard error in brackets. P = p-value of the difference between lines for each generation; ns = not significant; t (tendency) = P <0.10; * = P < 0.05; ** = P < 0.01; *** = P < 0.001.

|  | Unadjusted Model |  |  |  | Adjusted Model |  |  |  |
| --- | --- | --- | --- | --- | --- | --- | --- | --- |
|  | LRFI | HRFI | P |  | LRFI | HRFI | P |  |
| DAY |  |  |  |  |  |  |  |  |
| G0 | 1636 (77) | 1551 (77) | 0.4266 | ns | 1636 (42) | 1672 (42) | 0.5407 | ns |
| G1 | 1665 (16) | 1649 (16) | 0.4677 | ns | 1665 (9) | 1645 (8) | 0.0875 | t |
| G2 | 1560 (16) | 1597 (16) | 0.1055 | ns | 1618 (9) | 1647 (9) | 0.0196 | * |
| G3 | 1591 (16) | 1638 (17) | 0.0419 | * | 1598 (9) | 1600 (9) | 0.9201 | ns |
| G4 | 1546 (15) | 1611 (16) | 0.0035 | ** | 1616 (8) | 1593 (9) | 0.0554 | T |
| G5 | 1564 (15) | 1708 (16) | <.0001 | *** | 1621 (8) | 1618 (9) | 0.7821 | ns |
| G6 | 1573 (10) | 1724 (11) | <.0001 | *** | 1626 (6) | 1632 (6) | 0.4384 | ns |
| G7 | 1531 (11) | 1701 (12) | <.0001 | *** | 1634 (6) | 1610 (6) | 0.0100 | ** |
| G8 | 1619 (15) | 1727 (17) | <.0001 | *** | 1622 (8) | 1543 (9) | <.0001 | *** |
| G9 | 1522 (17) | 1672 (17) | <.0001 | *** | 1587 (10) | 1549 (10) | 0.0054 | ** |
| NIGHT |  |  |  |  |  |  |  |  |
| G0 | 424 (48) | 345 (48) | 0.2348 | ns | 424 (42) | 388 (42) | 0.5407 | ns |
| G1 | 395 (10) | 417 (10) | 0.1066 | ns | 395 (9) | 416 (8) | 0.0875 | t |
| G2 | 421 (10) | 395 (10) | 0.0654 | t | 442 (9) | 413 (9) | 0.0196 | * |
| G3 | 460 (10) | 475 (10) | 0.2901 | ns | 462 (9) | 461 (9) | 0.9201 | ns |
| G4 | 419 (10) | 474 (10) | 0.0001 | *** | 444 (8) | 468 (9) | 0.0554 | t |
| G5 | 419 (9) | 475 (10) | <.0001 | *** | 439 (8) | 443 (9) | 0.7821 | ns |
| G6 | 415 (6) | 461 (7) | <.0001 | *** | 434 (6) | 428 (6) | 0.4384 | ns |
| G7 | 390 (7) | 483 (7) | <.0001 | *** | 427 (6) | 450 (6) | 0.0100 | ** |
| G8 | 437 (9) | 584 (11) | <.0001 | *** | 438 (8) | 517 (9) | <.0001 | *** |
| G9 | 449 (11) | 556 (11) | <.0001 | *** | 473 (10) | 511 (10) | 0.0054 | ** |

### Distribution of average diurnal intake along selection

Table 3 reports the least-squares means for each line x generation combination for unadjusted and adjusted DFI during the peaks of feed consumption and between peaks.

**Table 3.** Average intake during first peak (06:00-10:00), second peak (15:00-21:00) and between peaks for each line (LRFI or HRFI) and generation (G0 to G9) combination (in g/period). Least squares of line x generation effect in a linear model with (adjusted) or without (unadjusted) average daily intake as covariate, standard error in brackets. P = p-value of the difference between lines for each generation; ns = not significant; t (tendency)= P <0.10; * = P < 0.05; ** = P < 0.01; *** = P < 0.001; **** = P < 0.0001.

|  | Unadjusted model |  |  |  | Adjusted model |  |  |  |
| --- | --- | --- | --- | --- | --- | --- | --- | --- |
|  | LRFI | HRFI | P |  | LRFI | HRFI | P |  |
| Peak 1 |  |  |  |  |  |  |  |  |
| G0 | 429 (42) | 327 (42) | 0.0830 | t | 429 (39) | 358 (39) | 0.1876 | ns |
| G1 | 397 (9) | 395 (8) | 0.8511 | ns | 397 (8) | 394 (8) | 0.7601 | ns |
| G2 | 397 (9) | 421 (9) | 0.0532 | t | 411 (8) | 433 (8) | 0.0546 | t |
| G3 | 368 (9) | 406 (9) | 0.0023 | ** | 370 (8) | 396 (8) | 0.0205 | * |
| G4 | 344 (8) | 381 (9) | 0.0019 | ** | 361 (8) | 377 (8) | 0.1630 | ns |
| G5 | 381 (8) | 385 (9) | 0.7623 | ns | 395 (8) | 362 (8) | 0.0035 | ** |
| G6 | 375 (6) | 407 (6) | 0.0002 | *** | 389 (5) | 384 (6) | 0.5425 | ns |
| G7 | 394 (6) | 407 (6) | 0.1344 | ns | 419 (6) | 384 (6) | <.0001 | **** |
| G8 | 434 (8) | 398 (9) | 0.0038 | ** | 435 (8) | 352 (9) | <.0001 | **** |
| G9 | 392 (10) | 366 (9) | 0.0539 | t | 408 (9) | 335 (9) | <.0001 | **** |
| Peak 2 |  |  |  |  |  |  |  |  |
| G0 | 734 (57) | 717 (57) | 0.8284 | ns | 734 (48) | 774 (48) | 0.5430 | ns |
| G1 | 890 (12) | 868 (11) | 0.1621 | ns | 890 (10) | 865 (10) | 0.0688 | t |
| G2 | 839 (12) | 794 (12) | 0.0082 | ** | 866 (10) | 818 (10) | 0.0007 | *** |
| G3 | 817 (12) | 792 (12) | 0.1357 | ns | 821 (10) | 774 (10) | 0.0010 | ** |
| G4 | 777 (11) | 777 (12) | 0.9756 | ns | 810 (10) | 768 (10) | 0.0022 | ** |
| G5 | 809 (11) | 823 (12) | 0.3924 | ns | 836 (9) | 779 (10) | 0.0001 | **** |
| G6 | 857 (8) | 820 (8) | 0.0011 | ** | 882 (6) | 777 (7) | <.0001 | **** |
| G7 | 809 (8) | 803 (9) | 0.6162 | ns | 858 (7) | 760 (7) | <.0001 | **** |
| G8 | 853 (11) | 814 (13) | 0.0203 | * | 855 (9) | 726 (11) | <.0001 | **** |
| G9 | 821 (13) | 821 (13) | 0.9776 | ns | 852 (11) | 763 (11) | <.0001 | **** |
| Inter- peaks |  |  |  |  |  |  |  |  |
| G0 | 531 (46) | 551 (46) | 0.7633 | ns | 531 (42) | 587 (42) | 0.3375 | ns |
| G1 | 458 (9) | 469 (9) | 0.4250 | ns | 458 (9) | 467 (8) | 0.4441 | ns |
| G2 | 413 (10) | 457 (10) | 0.0011 | ** | 431 (9) | 473 (9) | 0.0007 | *** |
| G3 | 489 (9) | 513 (10) | 0.0720 | t | 491 (9) | 502 (9) | 0.3767 | ns |
| G4 | 478 (9) | 517 (10) | 0.0035 | ** | 499 (8) | 512 (9) | 0.3074 | ns |
| G5 | 452 (9) | 570 (10) | <.0001 | **** | 470 (8) | 543 (9) | <.0001 | **** |
| G6 | 414 (6) | 568 (7) | <.0001 | **** | 430 (6) | 541 (6) | <.0001 | **** |
| G7 | 396 (7) | 560 (7) | <.0001 | **** | 427 (6) | 533 (6) | <.0001 | **** |
| G8 | 406 (9) | 585 (10) | <.0001 | **** | 407 (8) | 529 (9) | <.0001 | **** |
| G9 | 375 (10) | 560 (10) | <.0001 | **** | 394 (9) | 523 (9) | <.0001 | **** |

During peaks 1 and 2, according to unadjusted models, there were no consistent differences between the two RFI lines. On the opposite, when adjusted for DFI, there were higher intakes in the LRFI line at peaks: from generation G7 during peak 1 and from generation G2 during peak 2 onwards. The difference reached +80 g/day in the second peak in G9 for the LRFI pigs compared to the HRFI pigs, i.e., ∼10% of the intake of the period.

A different pattern emerged from the analysis of feed intake between peaks: the unadjusted model revealed significant differences between lines as early as generation G2, with the LRFI line ingesting less between peaks than the HRFI line (-185 g/day). After adjustment for DFI, this difference between lines persisted (P < 0.001) from generation G5 onwards, reaching -128 g/day in G9 for LRFI pigs as compared to HRFI pigs.

### Variance component estimates

The RFI had a heritability of 0.11 ± 0.02, and DFI 0.33 ± 0.04. The feeding rhythm traits were heritable (Tables 4 and 5). Heritability estimates were generally moderate for feeding pattern traits per visit (Table 4): they ranged from 0.45 ± 0.05 for feeding rate to 0.42 ± 0.05 for interval duration between visits.

**Table 4.** Heritability (h^2^) with standard errors in brackets for feeding pattern traits per visit.

| Traits | $h^2$ | $r_g$ | | | |
| --- | --- | --- | --- | --- | --- |
|  |  | RFI | Quantity fed per visit | Time spent per visit | Interval between visits |
| Quantity fed per visit | 0.44 (0.05) | -0.09 (0.10) |  |  |  |
| Time spent per visit | 0.43 (0.05) | -0.11 (0.10) | 0.91 (0.02) |  |  |
| Interval between visits | 0.42 (0.05) | -0.17 (0.10) | 0.99 (0.01) | 0.89 (0.02) |  |
| Feeding rate per visit | 0.45 (0.05) | -0.02 (0.10) | 0.23 (0.09) | -0.19 (0.09) | 0.23 (0.09) |

**Table 5.**
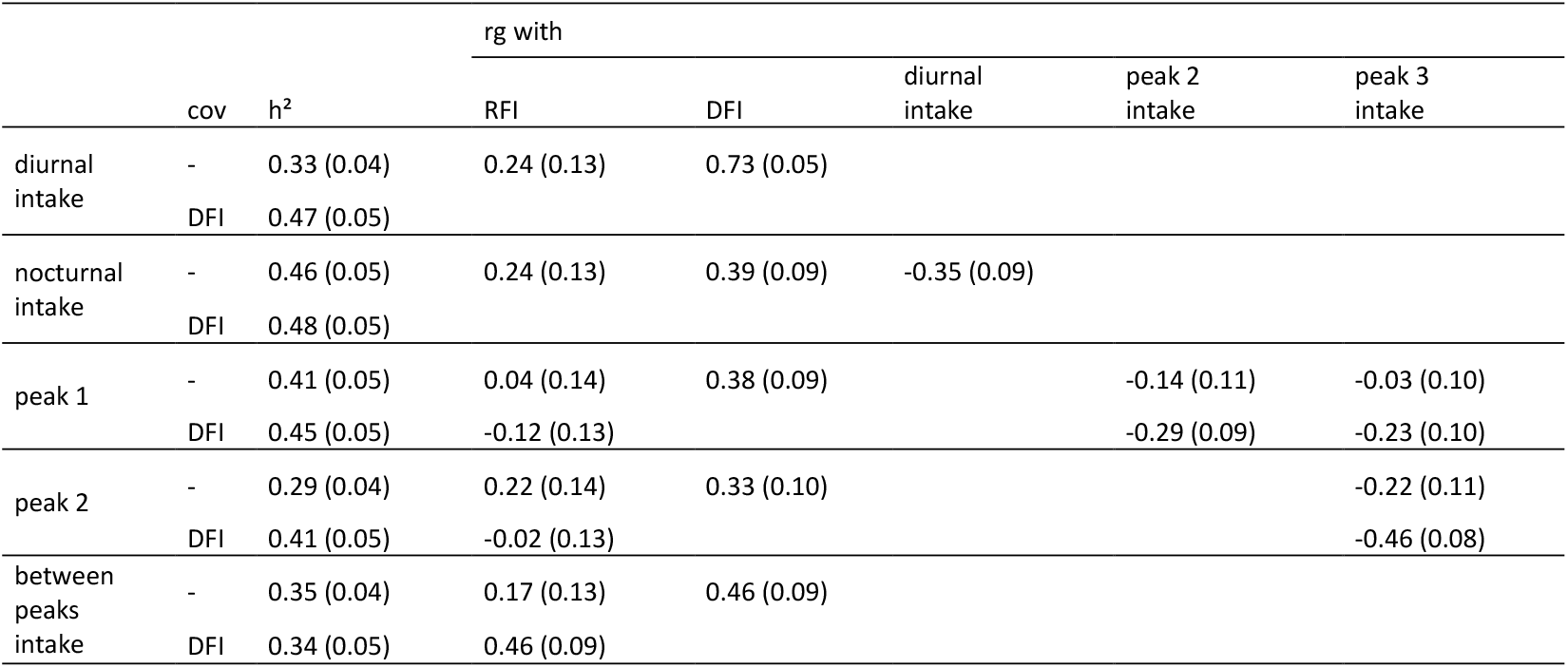
Variance components of the diurnal and nocturnal feed intakes (h^2^=heritability, rg = genetic correlation, standard error between parentheses), unadjusted for DFI (cov = -) or adjusted (cov = DFI)

For traits considered by periods (diurnal or nocturnal) (Table 5), heritability estimates ranged from 0.29 ± 0.04 to 0.47 ± 0.05 depending on the trait and whether or not it was adjusted to DFI. Genetic correlations with RFI were low (< 0.24 ± 0.13), except for DFI-adjusted inter-peak feed intake (0.46 ± 0.09). As expected, genetic correlations with DFI were higher than with RFI, between 0.33 ± 0.10 for the feed intake at peak 2 to 0.46 ± 0.09 for the feed intake between peaks.

Nocturnal feed intake was negatively correlated with diurnal feed intake (-0.35 ± 0.09). Feed intakes during and between peaks were negatively correlated when unadjusted for DFI. Correlation coefficients were greater when DFI adjusted values were considered (from -0.23 ± 0.10 for between intake at peak 2 and the between-peak period to -0.46 (0.08) for intakes between peak 2 and the between-peak period).

### Evolution of allele frequencies in clock genes during selection for RFI

Figure 2 summarizes the results of regression analyses on the SNPs allele frequencies applied to the 145,448 SNPs located in the gene coding sequences + 10^6^ bp around. A total of 3,497 SNPs were suggested at the chromosome-wide threshold (-log_10_(P-value > 6), among which 2,843 in HRFI line and 654 in LRFI line. These SNPs were located around *RORA, PER1, PER2, CRY2, ARNTL, CRY1* and *CLOCK* genes for HRFI line and around *NR1D1, PER1, PER2, ARNTL* genes in LRFI line. At the genome-wide level, in the HRFI line, changes in allele frequencies of SNPs within and around *ARNTL* and *CLOCK* genes were identified (p < 0.05 genome-wide). For *ARNTL*, 56 SNPs were detected, including 3 within the coding sequence, spanning from 81 kb upstream to 493kb downstream the gene, corresponding to average decreases of allele frequencies of 4.4% per generation (from ∼37% in G0 to 0 in G9). For the *CLOCK* gene, 334 SNPs were significant, spanning almost all the studied interval (from -937kb to +884kb to the coding sequence), including 10 SNPs within the coding sequence. They corresponded to SNPs in complete linkage disequilibrium, losing 1% minor allele frequency along the generations in the HRFI line, from 9% in G0 to 0 in G9. In the LRFI line, 16 SNPs were detected as having a genome-wide significant change in allele frequency between 665 and 403kb upstream the *NR1D1* gene (REVERBα). No significant SNP was detected jointly in the LRFI and HRFI lines.

**Figure 2.**
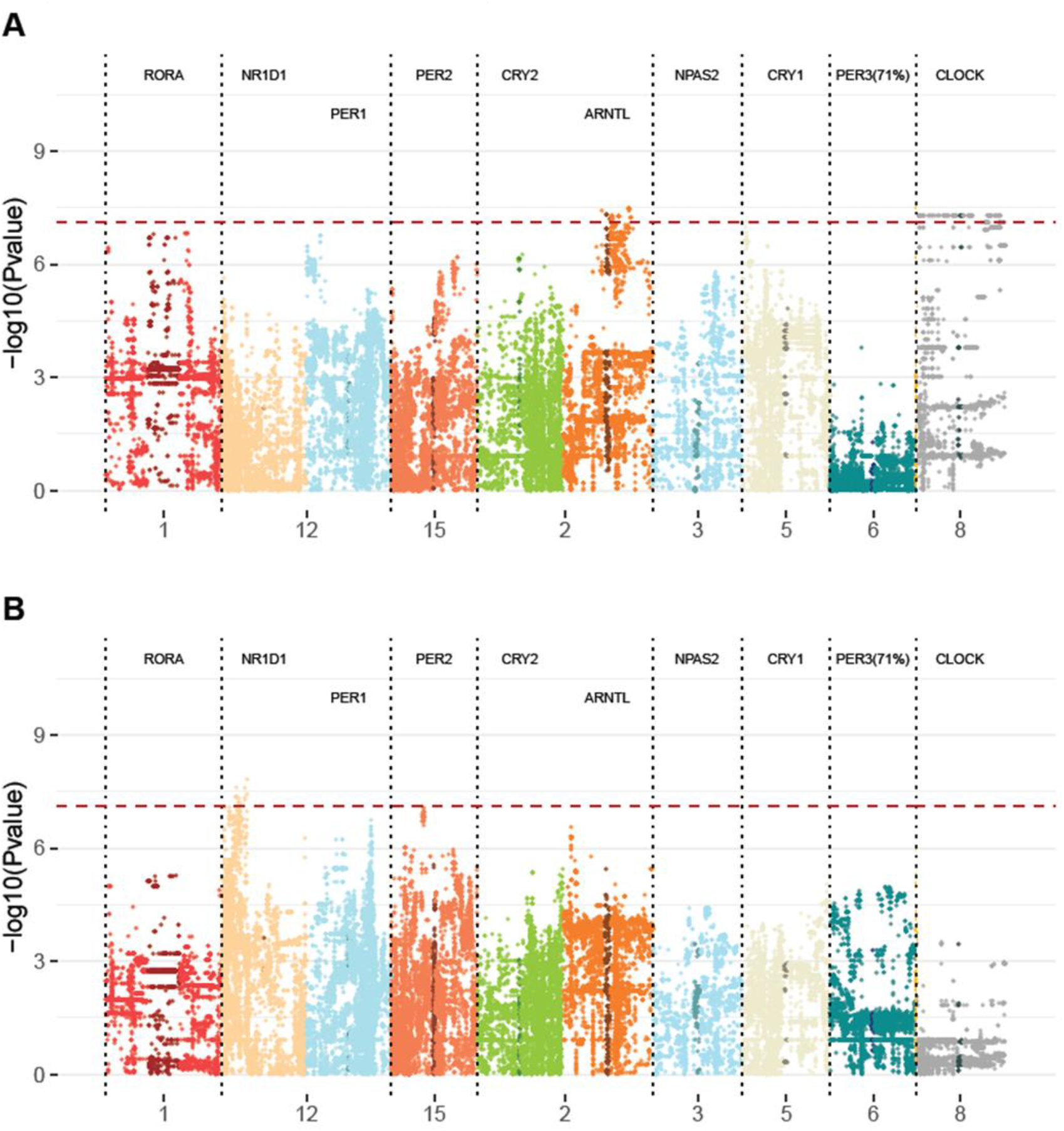
−log10(P value) of the regression of generation number on allele frequency for the 145,448 SNPs in the coding sequence of the core clock genes and 10^6^ bp around, in the HRFI (A) and LRFI (B) lines. Each point represents a SNP, positioned according to its position on the chromosome interval, ordered according to chromosome (SSC). SNPs in the coding sequence have a darker colour for each gene. The red dotted line represents the genome-wide threshold (-log10(P value) = 7.22).

**Figure 3.**
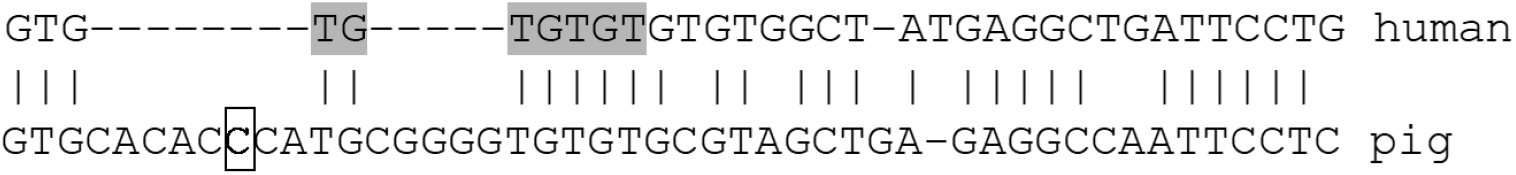
CRY2 3’UTR regions in human and pig. The has-miR-8485 miRNA binding region identified by Dfam database in the human sequence is highlighted in grey. The SNP identified in the pig sequence is the framed nucleotide.

### Putative functional effects of the SNPs

#### Variant Effect Predictor analysis

For the analysis of putative functional effects of the SNPs, we used the 145,448 SNPs of the dataset (clock genes coding sequence ± 10^6^ pb) and selected the 1,886 SNPs in common with the evolution of allele frequencies analysis. Using the Ensembl Variant Effect Predictor, we identified 256 SNPs with putative functional effects, 38 of which were located inside the coding sequence of *RORA, CRY2, CLOCK* and *ARNTL* genes, all of them in the HRFI line (Table 6) and 218 outside the clock gene coding sequence (59 for LRFI line, 159 for HRFI line, Table 7).

**Table 6.**
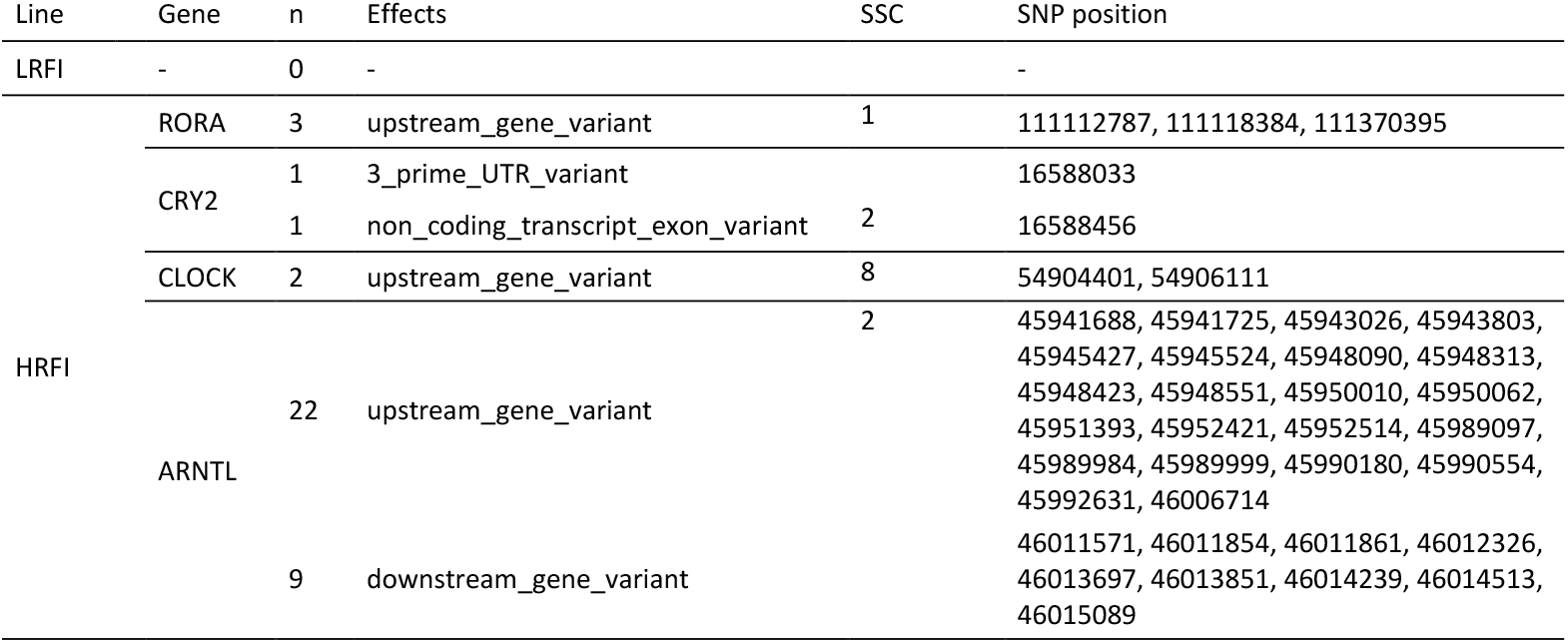
Characteristics of the SNPs identified inside the clock genes coding sequence, selected with the Ensembl Variant Effect Predictor for the following effects: 3_prime_UTR_variant, 5_prime_UTR_variant, downstream_gene_variant, missense_ variant, non_coding_transcript_exon_variant, non_coding_transcript_ variant, start_lost, stop_gained, stop_lost, stop_retained_variant, upstream_ gene_variant.

**Table 7.** Characteristics of the SNPs identified outside the clock genes coding sequence (+ 10^6^ bp around), selected with the Ensembl Variant Effect Predictor for the following effects: 3_prime_UTR_variant, 5_prime_UTR_variant, downstream_ gene_variant, missense_variant, non_coding_transcript_exon_ variant, non_ coding_transcript_variant, start_lost, stop_gained, stop_lost, stop_retained_variant, upstream_ gene_variant.

| Line | Effect | SSC | n | SNPs | Gene |
| --- | --- | --- | --- | --- | --- |
| LRFI | missense variant | 12 | 16 | 21641370<br>21600701<br>21689439, 21689449, 21689451, 21689473, 21690361, 21691270, 21691314, 21691623, 21692765<br>21677265<br>21669015<br>21285791<br>21422455, 21439835 | KRT10<br>KRT12<br>KRT26<br>KRT27<br>KRT28<br>KRTAP29-1<br>no annotation |
|  | non coding transcript exon variant | 12 | 43 | 21689312, 21689586, 21689650<br>21679474<br>21611041, 21611197, 21611244, 21611346, 21611441, 21611447, 21611669, 21611743, 21611745, 21611749, 21611995, 21612015, 21613105, 21613126, 21670735, 21670755, 21670762, 21670806, 21670937, 21670955, 21671386, 21671553, 21671593, 21671610, 21671650, 21671707, 21671717, 21671733, 21672047, 21672048, 21672058, 21672069, 21672129, 21672190, 21672297, 21672312, 21672952, 21673660, 21759695 | KRT26<br>KRT27<br>long non coding RNA |
| HRFI | missense variant | 2 | 2 | 16563362<br>46337272 | FREY1<br>RASSF10 |
|  |  | 8 | 18 | 55609178, 55611921, 55619533, 55619593, 55642273<br>55763830<br>55562997, 55598132<br>55868067, 55901550, 55901580<br>55759547, 55759687<br>55787351, 55787473, 55787521, 55787612, 55811369 | AASDH<br>ARL9<br>CRACD<br>HOPX<br>SRP72<br>THEGL |
|  |  | 12 | 1 | 152684513 | no annotation |
|  | non coding transcript exon variant | 2 | 2 | 46453151<br>46735824 | TEAD<br>long non coding RNA |
|  |  | 5 | 7 | 12392661, 12395439, 12395694, 12400668, 12401018, 12401325, 12401993 | no gene |
|  |  | 8 | 128 | 55862284, 55862358, 55862371, 55862493, 55862811, 55867347, 55867627<br>54231930, 55058364, 55058382, 55061173, 55063034, 55063267, 55063894, 55064101, 55065022, 55065600, 55066759, 55067663, 55068258, 55069790, 55071273, 55071707, 55072891, 55072953, 55078235, 55078246, 55078453, 55078811, 55079711, 55079712, 55081112, 55472845, 55473186, 55473417, 55473547, 55473844, 55473875, 55474486, 55474605, 55475130, 55475335, 55475701, 55476634, 55476835, 55477024, 55477335, 55477509, 55477517, 55478140, 55478333, 55478599, 55478603, 55478878, 55480275, 55480368, 55480520, 55480542, 55480578, 55480601, 55480671, 55481033, 55485903, 55486165, 55486449, 55486979, 55487157, 55487158, 55487186, 55487563, 55487626, 55487854, 55488055, 55488062, 55488167, 55488495, 55488500, 55489174, 55489421, 55489571, 55489587, 55489592, 55489977, 55490143, 55490360, 55499839, 55499955, 55500255, 55500403, 55500436, 55500637, 55501849, 55502460, 55502475, 55502847, 55502942, 55503094, 55504364, 55504908, 55506407, 55507433, 55507627, 55507685, 55507741, 55508838, 55509509, 55510492, 55511433, 55512162, 55512208, 55512268, 55515033, 55515338, 55516652, 55516688, 55517044, 55517405, 55517518, 55518513, 55518715, 55519296, 55519408, 55519659<br>55531525, 55601118, 55601428, 55602744, 55603322 | HOPX<br>long non coding RNA<br>no gene |
|  | Stop gained | 8 | 1 | 55867753 | HOPX |

#### Presence of SNPs in miRNA binding regions

*CRY2* was the only clock gene showing a SNP in the 3’UTR region (position 16,588,033), where a potential miRNA binding region was located. Using the miRDB target prediction database we found that hsa-miR-8485 miRNA binds the 3’UTR region of human *CRY2* (Figure 4). However, the SNP identified in the pig *CRY2* sequence was located in a region that did not match the human sequence.

**Figure 4.**
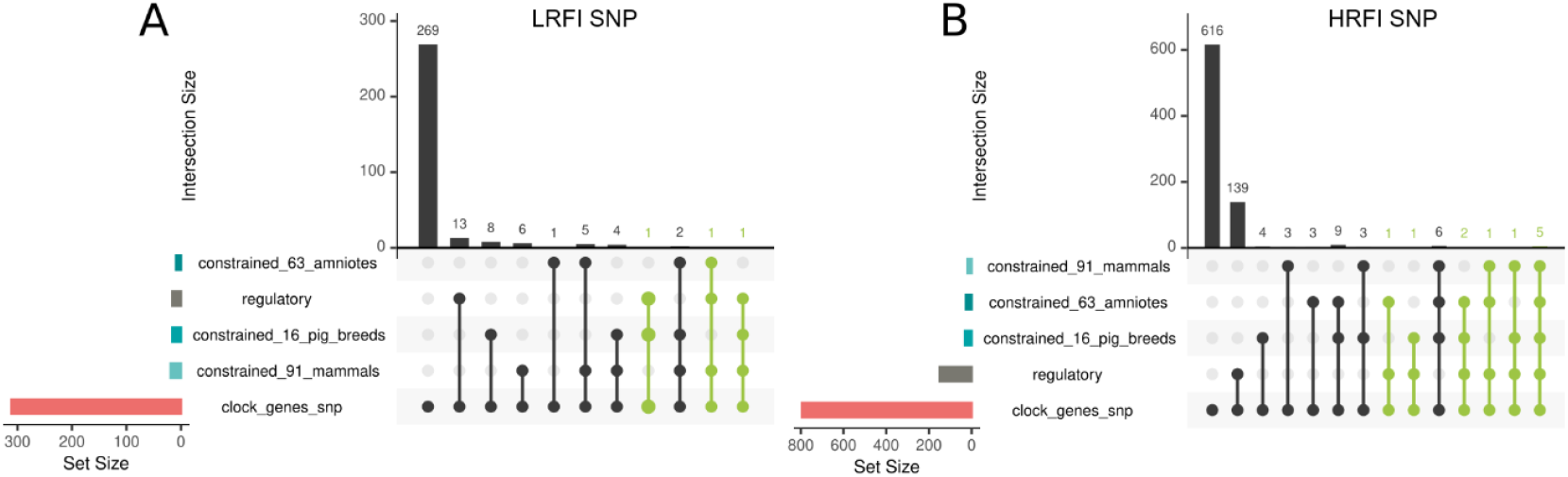
Number of significant SNPs from clock genes (coding sequence ± 10^6^ bp, clock_genes_snp) intersecting conserved or regulatory genomic regions: mammalian (constrained_91_mamals), amniote (constrained_63_amniotes), pig breeds (constrained_16_pig_breeds) and pig regulatory regions, for LRFI (A) and HRFI lines (B).

#### Presence of SNPs in Transposable Elements DNA sequences

Using the Dfam database we found that 7 SNPs of the *ARNTL* gene are located in Transposable Elements: 2 SNPs (45,950,010 and 45,950,062) in the MIRb (Mammalian-wide Interspersed Repeat - variant b) TE, 1 SNP (45,952,514) in the MIRc (Mammalian-wide Interspersed Repeat - variant c) TE, 2 SNPs (45,943,026 and 45,943,803) in the L2 (LINE2) retrotransposon and 2 SNPs (45,945,427 and 45,945,524) in the *ORF2* from L1 retrotransposon, L1M5_orf2 subfamily.

#### SNPs conservation level and localization to regulatory regions

Among the SNPs showing a suggestive change in allele frequency during selection (-log_10_(P-value>6) from pig core clock genes (coding sequence + 10^6^ bp around), we identified the SNPs that are in conserved regions at three phylogenic levels (mammalian, amniote, pig breeds), or localized in regulatory regions for each line (Figure 5).

For the LRFI line, 3 SNP were both in a regulatory region and in a conserved region (in at least one of the three level of conservation used). Two of these three SNP were located in a putative transcription factor (TF) binding site as identified by mapping JASPAR TF motifs on the pig reference genome: SNP 121,285,791 was in a MEIS1 motif, and 121,422,455 in a ZFP57 and SP5 motif (Table 8).

**Table 8.**
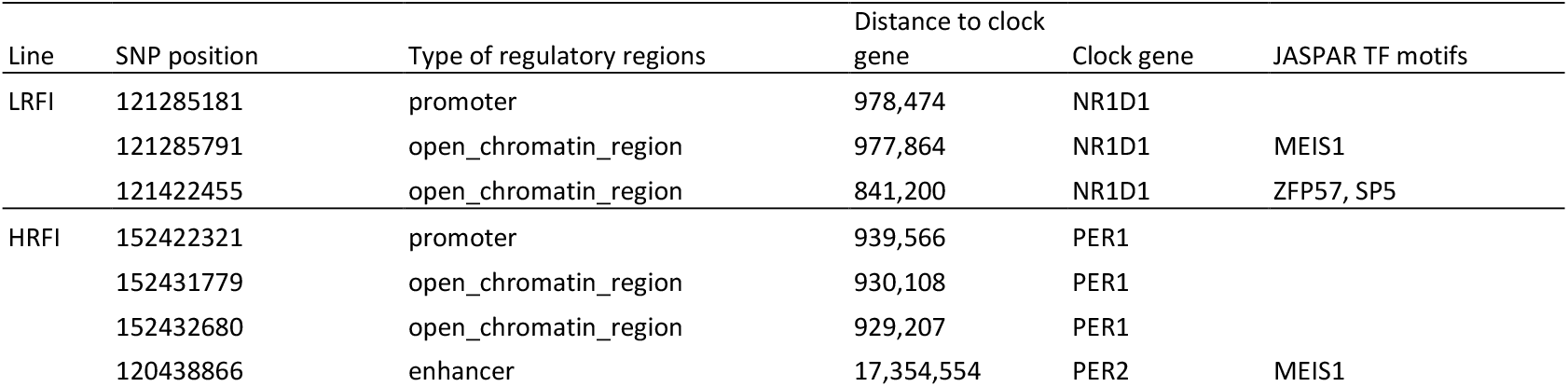
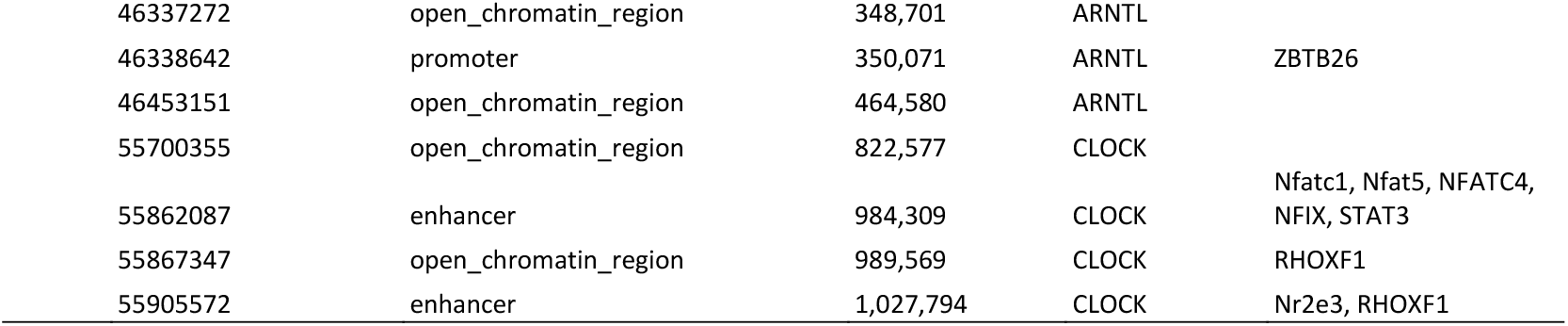
Significant SNP responding to selection near clock genes, falling in conserved and regulatory genomic regions. The last column indicates JASPAR transcription factor motif that are predicted to be altered by the SNP.

| Line | SNP position | Type of regulatory regions | Distance to clock gene | Clock gene | JASPAR TF motifs |
| --- | --- | --- | --- | --- | --- |
| LRFI | 121285181 | promoter | 978,474 | NR1D1 |  |
|  | 121285791 | open_chromatin_region | 977,864 | NR1D1 | MEIS1 |
|  | 121422455 | open_chromatin_region | 841,200 | NR1D1 | ZFP57, SP5 |
| HRFI | 152422321 | promoter | 939,566 | PER1 |  |
|  | 152431779 | open_chromatin_region | 930,108 | PER1 |  |
|  | 152432680 | open_chromatin_region | 929,207 | PER1 |  |
|  | 120438866 | enhancer | 17,354,554 | PER2 | MEIS1 |

|  |  |  |  |  |
| --- | --- | --- | --- | --- |
| 46337272 | open_chromatin_region | 348,701 | ARNTL |  |
| 46338642 | promoter | 350,071 | ARNTL | ZBTB26 |
| 46453151 | open_chromatin_region | 464,580 | ARNTL |  |
| 55700355 | open_chromatin_region | 822,577 | CLOCK |  |
| 55862087 | enhancer | 984,309 | CLOCK | Nfatc1, Nfat5, NFATC4, NFIX, STAT3 |
| 55867347 | open_chromatin_region | 989,569 | CLOCK | RHOXF1 |
| 55905572 | enhancer | 1,027,794 | CLOCK | Nr2e3, RHOXF1 |

For the HRFI line, 11 SNP were located both in a regulatory region and in a conserved region. Five of these SNP were located in a putative transcription factor site (Table 8), involving motifs for MEIS1, ZBTB26, RHOXF1, NFIX, Nfatc1, Nfat5, NFATC4, STAT3, RHOXF1, and NR2E3.

## Discussion

Improving feed efficiency in growing pigs is a priority because of the high costs of feed and the correlated benefits for reducing environmental impacts. The influence of circadian rhythms on feed efficiency has not been studied in pigs. In this study, we described the feeding rhythms and the polymorphisms in core clock genes in relation with the variability of feed efficiency, by using two pig lines genetically selected for contrasting RFI along ten generations.

By analyzing 6,494,097 visits to automatic concentrate dispensers across a population of 3,824 pigs, we identified distinct differences in feeding behavior between low and high RFI pig lines. When averaged along the generations of selection, we found that the LRFI pigs eat less frequently, with larger meals and of a longer duration; they also exhibited a higher feeding rate at each visit when compared to the HRFI pigs. These findings corroborate earlier studies on the same lines, conducted at an earlier generation of selection (G7) using video scan samplings along the growing and finishing period (Meunier-Salaün et al., 2014). This first study suggested that the energy cost related to the standing posture associated to visits at the feed dispensers may be part of the explanation of the difference in the feed efficiency between LRFI and HRFI pigs. Importantly, thanks to the data obtained from automatic feed dispensers, we further show that LRFI pigs displayed a distinct feeding pattern during the day, which was characterized by two peaks of feed intake: the first peak occurring around 8:00 and a second, which was higher, at around 17:00. This confirms previous studies in growing pigs showing that spontaneous feeding rhythm is characterized by two consumption’s peaks, one in the morning and one in the afternoon (Andretta et al., 2016; Renaudeau, 2009), and that peak intake in the late afternoon is generally greater than that in the morning (Boumans et al., 2017; Poullet et al., 2022). In contrast, the HRFI pigs ate more evenly throughout the day, particularly between the two peaks, which results in a smoother feeding pattern during the day. These differences between the two RFI lines were accentuated as the number of generations of selection increased. This suggests that different genetic bases associated with feeding behavior have been retained in each line. Another study examining the individual eating behaviors of 110 commercial pigs over 83 days revealed substantial variations in eating patterns: 57% of the pigs exhibited circadian feeding rhythms with two peaks, while the remaining pigs displayed more irregular feeding patterns with intermittent periods of non-rhythmic behavior (Bus et al., 2023). Factors such as age, competition at the feeder, and individual personality were suggested as influencing circadian feeding behavior in that study (Bus et al., 2023). Our data revealed that, in addition to these environmental factors, different genetic polymorphisms may also contribute to the observed differences across successive generations of selection.

The smoothing of feeding behavior peaks along the day which was observed in the HRFI pigs may indicate either the loss of circadian rhythms or a greater variability in feeding behavior from day to day in this line. Importantly, we observed that LRFI pigs consumed more feed during the diurnal period and less at night when compared to HRFI pigs, starting from generation G7 onwards, relatively to line differences in average daily feed intake. The nychthemeral profile of behaviors in Large White pigs indicates higher active behaviors during the diurnal period than during the night (Meunier-Salaün et al., 2014). Irregular or erratic meal patterns, increased frequency of eating occasions, nocturnal calorie intake, and prolonged eating durations are all factors that can contribute to poorer metabolic health (Flanagan et al., 2021). The frequency and size of the meals can affect the digestibility of nutrients by modulating gastrointestinal tract conditions (pH and transit time), metabolites (glucose and short-chain fatty acids) and hormones (Chassé et al., 2021). For instance, when young growing crossbred pigs were offered a same amount of feed either in 2 or 12 meals per day during a 3-wk interventional period, the more frequent daily meal intake decreased the conversion of feed into weight gain (van Erp et al., 2020). Also, when intake is shifted towards night hours, fat deposition is increased in pigs (van Erp et al., 2020). Therefore, the progressive shift in feeding behavior for the HRFI pigs having a marked nibbling behavior rather than a meal behavior can be one of the biological bases of poorer use of nutrients, and thus of deteriorated feed efficiency at the animal level in this line.

In this study, the genetic analyses revealed that traits related to feed intake during the two main diurnal peaks, as well as intake between these peaks, were heritable, with heritability estimates (h^2^) ranging from 0.30 to 0.40. Heritability of circadian rhythms has been previously reported in both mice and humans (Vitaterna et al., 2019). The genetic correlation between feed intake and RFI was generally low, except for intake between the peaks where the correlation was moderate (r = 0.46). A moderate to high genetic correlation was also observed with daily feed intake. these genetic links sustained the possibility that indirect selection could occur for circadian rhythm in response to selection for RFI. Since circadian rhythms are primarily regulated by core clock genes, we investigated the evolution of allele frequencies in DNA sequences surrounding these genes along generations of selection. By searching for SNPs in the coding sequences and 10^6^ base pairs upstream and downstream of each core clock gene, we detected a higher number of SNPs with significant changes in frequencies in the HRFI line compared to the LRFI line (2,843 SNPs in HRFI versus 654 in LRFI, respectively). The most significant SNP, located in the coding and flanking sequences of *ARNTL* and *CLOCK* genes in the HRFI line, suggests that haplotypes in these genes have evolved during selection for RFI, particularly in this line. At the chromosome-wide level, SNPs located within the coding sequences were found exclusively in the HRFI line, and were identified in the genes *RORA, CRY2, CLOCK*, and *ARNTL*. Notably, *CRY2* displayed a SNP in its 3’ untranslated region (UTR). However, no miRNA binding site was detected. For the other genes, the SNPs were located either upstream or downstream of the coding sequence. To further assess the functional significance of these SNPs, we examined whether their locations overlapped with transposable elements (TEs) or conserved regulatory regions. Among these, only SNPs in the *ARNTL* coding region were found to be located within a transposable element and 2 SNPs in *CRY2* and 4 SNPs in *ARNTL* coding regions were located in regulatory regions, which sustains a potential functionality associated with these frequency changes.

Genes *ARNTL* and *CLOCK* are coding for transcription factors that form heterodimers to regulate the oscillatory expression of rhythmic genes across cells in various organs of the body. In both mice and baboons, it has been estimated that 20% to 30% of genes exhibit rhythmic expression in organs such as the liver, adipose tissue, and muscle (Mure et al., 2018; Zhang et al., 2014). This suggests that polymorphisms in these two genes could have significant effects on metabolism, and thus influence RFI. Moreover, *CRY2* is a target gene of the ARNTL-CLOCK complex and forms a heterodimer with PER proteins, which in turn represses their own transcription through a feedback loop with the CLOCK-ARNTL complex. Another key target gene of ARNTL-CLOCK is *NR1D1*, which negatively regulates *ARNTL*. Mutations in these genes or disruptions in their regulatory mechanisms have been reported to cause changes in sleep patterns and energy metabolism in several species, including mice (Laposky et al., 2005; Naylor et al., 2000) and humans (Mishima et al., 2005), with broader implications for metabolic health (see Škrlec et al., 2021 for review). In the current study, we observed a significant accumulation of SNPs changing frequency within the coding sequence of the *ARNTL* gene in the HRFI line. While the functional consequences of these SNPs remain unclear, their location within transposable elements or potential regulatory regions are intriguing new avenues for research. As for the SNPs outside the coding regions of the core clock genes, those that are conserved across species— whether located within regulatory regions or not—warrant further investigation to determine their potential functional effects.

## Conclusion

This study highlights the important role of daily feeding patterns in the variation of feed efficiency in growing pigs. We identified distinct feeding behaviors between low and high RFI lines, with LRFI pigs exhibiting a more structured feeding pattern with two peaks, potentially reflecting more efficient energy use. Genetic analyses revealed that feeding behavior, particularly intake between the two peaks, is heritable and genetically correlated with daily feed intake, suggesting a genetic basis for the observed differences in feeding patterns. Further investigation of SNPs in core clock genes, such as *ARNTL, CLOCK, CRY2*, and *RORA*, particularly in regulatory regions and transposable elements, could provide insights into the genetic mechanisms underlying these feeding behaviors and their potential role in improving feed efficiency. These findings open perspectives for breeding strategies aimed at enhancing metabolic efficiency in pigs, which is critical for reducing feed costs and improving environmental sustainability in agriculture.

## Acknowledgements

We would like to thank Yann Labrune and Marc Teissier for their contribution to the molecular data analysis and the staff of INRAE (Rennes and Toulouse, France), in particular all the animal keepers and all the laboratory technicians for their collaboration during the experiment.

## Data availability

Data and script are available online: https://doi.org/10.57745/LN1IZE.

## References

Aliakbari, A., Delpuech, E., Labrune, Y., Riquet, J., Gilbert, H., 2020. The impact of training on data from genetically-related lines on the accuracy of genomic predictions for feed efficiency traits in pigs. Genet Sel Evol 52, 57. 10.1186/s12711-020-00576-0

Andretta, I., Pomar, C., Kipper, M., Hauschild, L., Rivest, J., 2016. Feeding behavior of growing-finishing pigs reared under precision feeding strategies. J Anim Sci 94, 3042–3050. 10.2527/jas.2016-0392

Boege, H.L., Bhatti, M.Z., St-Onge, M.-P., 2021. Circadian rhythms and meal timing: impact on energy balance and body weight. Current Opinion in Biotechnology 70, 1–6. 10.1016/j.copbio.2020.08.009

Boumans, I.J.M.M., de Boer, I.J.M., Hofstede, G.J., la Fleur, S.E., Bokkers, E.A.M., 2017. The importance of hormonal circadian rhythms in daily feeding patterns: An illustration with simulated pigs. Hormones and Behavior 93, 82–93. 10.1016/j.yhbeh.2017.05.003

Bus, J.D., Boumans, I.J.M.M., Engel, J., Te Beest, D.E., Webb, L.E., Bokkers, E.A.M., 2023. Circadian rhythms and diurnal patterns in the feed intake behaviour of growing-finishing pigs. Sci Rep 13, 16021. 10.1038/s41598-023-42612-1

Castro-Mondragon, J.A., Riudavets-Puig, R., Rauluseviciute, I., Berhanu Lemma, R., Turchi, L., Blanc-Mathieu, R., Lucas, J., Boddie, P., Khan, A., Manosalva Pérez, N., Fornes, O., Leung, T.Y., Aguirre, A., Hammal, F., Schmelter, D., Baranasic, D., Ballester, B., Sandelin, A., Lenhard, B., Vandepoele, K., Wasserman, W.W., Parcy, F., Mathelier, A., 2022. JASPAR 2022: the 9th release of the open-access database of transcription factor binding profiles. Nucleic Acids Research 50, D165–D173. 10.1093/nar/gkab1113

Chang, C.C., Chow, C.C., Tellier, L.C., Vattikuti, S., Purcell, S.M., Lee, J.J., 2015. Second-generation PLINK: rising to the challenge of larger and richer datasets. Gigascience 4, 7. 10.1186/s13742-015-0047-8

Chassé, É., Guay, F., Bach Knudsen, K.E., Zijlstra, R.T., Létourneau-Montminy, M.-P., 2021. Toward Precise Nutrient Value of Feed in Growing Pigs: Effect of Meal Size, Frequency and Dietary Fibre on Nutrient Utilisation. Animals (Basel) 11, 2598. 10.3390/ani11092598

Chen, Y., Wang, X., 2020. miRDB: an online database for prediction of functional microRNA targets. Nucleic Acids Res 48, D127–D131. 10.1093/nar/gkz757

Conway, J.R., Lex, A., Gehlenborg, N., 2017. UpSetR: an R package for the visualization of intersecting sets and their properties. Bioinformatics 33, 2938–2940. 10.1093/bioinformatics/btx364

Delpuech, E., Aliakbari, A., Labrune, Y., Fève, K., Billon, Y., Gilbert, H., Riquet, J., 2021. Identification of genomic regions affecting production traits in pigs divergently selected for feed efficiency. Genet Sel Evol 53, 49. 10.1186/s12711-021-00642-1

Flanagan, A., Bechtold, D.A., Pot, G.K., Johnston, J.D., 2021. Chrono-nutrition: From molecular and neuronal mechanisms to human epidemiology and timed feeding patterns. Journal of Neurochemistry 157, 53–72. 10.1111/jnc.15246

Fox, J., Weisberg, S., 2019. An R Companion to Applied Regression, Sage, Third Edition, Thousand Oaks CA.

Gao, X., 2011. Multiple testing corrections for imputed SNPs. Genet Epidemiol 35, 154–158. 10.1002/gepi.20563

Gilbert, H., Bidanel, J.-P., Gruand, J., Caritez, J.-C., Billon, Y., Guillouet, P., Lagant, H., Noblet, J., Sellier, P., 2007. Genetic parameters for residual feed intake in growing pigs, with emphasis on genetic relationships with carcass and meat quality traits. J Anim Sci 85, 3182–3188. 10.2527/jas.2006-590

Gilbert, H., Billon, Y., Brossard, L., Faure, J., Gatellier, P., Gondret, F., Labussière, E., Lebret, B., Lefaucheur, L., Le Floch, N., Louveau, I., Merlot, E., Meunier-Salaün, M.-C., Montagne, L., Mormede, P., Renaudeau, D., Riquet, J., Rogel-Gaillard, C., van Milgen, J., Vincent, A., Noblet, J., 2017. Review: divergent selection for residual feed intake in the growing pig. Animal 11, 1427–1439. 10.1017/S175173111600286X

Gilmour, A.R., 2019. Average information residual maximum likelihood in practice. J Anim Breed Genet 136, 262–272. 10.1111/jbg.12398

Gondret, F., Vincent, A., Houée-Bigot, M., Siegel, A., Lagarrigue, S., Causeur, D., Gilbert, H., Louveau, I., 2017. A transcriptome multi-tissue analysis identifies biological pathways and genes associated with variations in feed efficiency of growing pigs. BMC Genomics 18, 244. 10.1186/s12864-017-3639-0

Harrison, P.W., Amode, M.R., Austine-Orimoloye, O., Azov, A.G., Barba, M., Barnes, I., Becker, A., Bennett, R., Berry, A., Bhai, J., Bhurji, S.K., Boddu, S., Branco Lins, P.R., Brooks, L., Ramaraju, S.B., Campbell, L.I., Martinez, M.C., Charkhchi, M., Chougule, K., Cockburn, A., Davidson, C., De Silva, N.H., Dodiya, K., Donaldson, S., El Houdaigui, B., Naboulsi, T.E., Fatima, R., Giron, C.G., Genez, T., Grigoriadis, D., Ghattaoraya, G.S., Martinez, J.G., Gurbich, T.A., Hardy, M., Hollis, Z., Hourlier, T., Hunt, T., Kay, M., Kaykala, V., Le, T., Lemos, D., Lodha, D., Marques-Coelho, D., Maslen, G., Merino, G.A., Mirabueno, L.P., Mushtaq, A., Hossain, S.N., Ogeh, D.N., Sakthivel, M.P., Parker, A., Perry, M., Piližota, I., Poppleton, D., Prosovetskaia, I., Raj, S., Pérez-Silva, J.G., Salam, A.I.A., Saraf, S., Saraiva-Agostinho, N., Sheppard, D., Sinha, S., Sipos, B., Sitnik, V., Stark, W., Steed, E., Suner, M.-M., Surapaneni, L., Sutinen, K., Tricomi, F.F., Urbina-Gómez, D., Veidenberg, A., Walsh, T.A., Ware, D., Wass, E., Willhoft, N.L., Allen, J., Alvarez-Jarreta, J., Chakiachvili, M., Flint, B., Giorgetti, S., Haggerty, L., Ilsley, G.R., Keatley, J., Loveland, J.E., Moore, B., Mudge, J.M., Naamati, G., Tate, J., Trevanion, S.J., Winterbottom, A., Frankish, A., Hunt, S.E., Cunningham, F., Dyer, S., Finn, R.D., Martin, F.J., Yates, A.D., 2024. Ensembl 2024. Nucleic Acids Res 52, D891–D899. 10.1093/nar/gkad1049

Kuznetsova, A., Brockhoff, P.B., Christensen, R.H.B., 2017. lmerTest Package: Tests in Linear Mixed Effects Models. Journal of Statistical Software 82, 1–26. 10.18637/jss.v082.i13

Laposky, A., Easton, A., Dugovic, C., Walisser, J., Bradfield, C., Turek, F., 2005. Deletion of the Mammalian Circadian Clock Gene BMAL1/Mop3 Alters Baseline Sleep Architecture and the Response to Sleep Deprivation. Sleep 28, 395–410. 10.1093/sleep/28.4.395

Manoogian, E.N.C., Zadourian, A., Lo, H.C., Gutierrez, N.R., Shoghi, A., Rosander, A., Pazargadi, A., Ormiston, C.K., Wang, X., Sui, J., Hou, Z., Fleischer, J.G., Golshan, S., Taub, P.R., Panda, S., 2022. Feasibility of time-restricted eating and impacts on cardiometabolic health in 24-h shift workers: The Healthy Heroes randomized control trial. Cell Metab 34, 1442-1456.e7. 10.1016/j.cmet.2022.08.018

McLaren, W., Gil, L., Hunt, S.E., Riat, H.S., Ritchie, G.R.S., Thormann, A., Flicek, P., Cunningham, F., 2016. The Ensembl Variant Effect Predictor. Genome Biol 17, 122. 10.1186/s13059-016-0974-4

Meunier-Salaün, M.C., Guérin, C., Billon, Y., Sellier, P., Noblet, J., Gilbert, H., 2014. Divergent selection for residual feed intake in group-housed growing pigs: characteristics of physical and behavioural activity according to line and sex. Animal 8, 1898–1906. 10.1017/S1751731114001839

Mishima, K., Tozawa, T., Satoh, K., Saitoh, H., Mishima, Y., 2005. The 3111T/C polymorphism of hClock is associated with evening preference and delayed sleep timing in a Japanese population sample. American J of Med Genetics Pt B 133B, 101–104. 10.1002/ajmg.b.30110

Mure, L.S., Le, H.D., Benegiamo, G., Chang, M.W., Rios, L., Jillani, N., Ngotho, M., Kariuki, T., Dkhissi-Benyahya, O., Cooper, H.M., Panda, S., 2018. Diurnal transcriptome atlas of a primate across major neural and peripheral tissues. Science 359, eaao0318. 10.1126/science.aao0318

Naylor, E., Bergmann, B.M., Krauski, K., Zee, P.C., Takahashi, J.S., Vitaterna, M.H., Turek, F.W., 2000. The Circadian Clock Mutation Alters Sleep Homeostasis in the Mouse. J. Neurosci. 20, 8138–8143. 10.1523/JNEUROSCI.20-21-08138.2000

Nielsen, B.L., Lawrence, A.B., Whittemore, C.T., 1995. Effect of group size on feeding behaviour, social behaviour, and performance of growing pigs using single-space feeders. Livestock Production Science 44, 73–85. 10.1016/0301-6226(95)00060-X

Noblet, J., Karege, C., Dubois, S., van Milgen, J., 1999. Metabolic utilization of energy and maintenance requirements in growing pigs: effects of sex and genotype. J Anim Sci 77, 1208– 1216. 10.2527/1999.7751208x

Panda, S., 2016. Circadian physiology of metabolism. Science 354, 1008–1015. 10.1126/science.aah4967

Potter, G.D.M., Skene, D.J., Arendt, J., Cade, J.E., Grant, P.J., Hardie, L.J., 2016. Circadian Rhythm and Sleep Disruption: Causes, Metabolic Consequences, and Countermeasures. Endocr Rev 37, 584–608. 10.1210/er.2016-1083

Poullet, N., Rauw, W.M., Renaudeau, D., Riquet, J., Giorgi, M., Billon, Y., Gilbert, H., Gourdine, J.-L., 2022. Plasticity of feeding behaviour traits in response to production environment (temperate vs. tropical) in group-housed growing pigs. Sci Rep 12, 847. 10.1038/s41598-021-04752-0

Quinlan, A.R., Hall, I.M., 2010. BEDTools: a flexible suite of utilities for comparing genomic features. Bioinformatics 26, 841–842. 10.1093/bioinformatics/btq033

R Core Team, 2021. R: A language and environment for statistical computing. R Foundation for Statistical Computing, Vienna, Austria. https://www.R-project.org/.

Renaudeau, D., 2009. Effect of housing conditions (clean vs. dirty) on growth performance and feeding behavior in growing pigs in a tropical climate. Trop Anim Health Prod 41, 559–563. 10.1007/s11250-008-9223-5

Sargolzaei, M., Chesnais, J.P., Schenkel, F.S., 2014. A new approach for efficient genotype imputation using information from relatives. BMC Genomics 15, 478. 10.1186/1471-2164-15-478

Škrlec, I., Talapko, J., Džijan, S., Cesar, V., Lazić, N., Lepeduš, H., 2021. The Association between Circadian Clock Gene Polymorphisms and Metabolic Syndrome: A Systematic Review and Meta-Analysis. Biology 11, 20. 10.3390/biology11010020

Storer, J., Hubley, R., Rosen, J., Wheeler, T.J., Smit, A.F., 2021. The Dfam community resource of transposable element families, sequence models, and genome annotations. Mob DNA 12, 2. 10.1186/s13100-020-00230-y

Takahashi, J.S., Hong, H.-K., Ko, C.H., McDearmon, E.L., 2008. The genetics of mammalian circadian order and disorder: implications for physiology and disease. Nat Rev Genet 9, 764– 775. 10.1038/nrg2430

Tan, G., Lenhard, B., 2016. TFBSTools: an R/bioconductor package for transcription factor binding site analysis. Bioinformatics 32, 1555–1556. 10.1093/bioinformatics/btw024

van Erp, R.J.J., de Vries, S., van Kempen, T.A.T.G., Den Hartog, L.A., Gerrits, W.J.J., 2020. Circadian misalignment imposed by nocturnal feeding tends to increase fat deposition in pigs. Br J Nutr 123, 529–536. 10.1017/S0007114519003052

Vitaterna, M.H., Shimomura, K., Jiang, P., 2019. Genetics of Circadian Rhythms. Neurologic Clinics 37, 487–504. 10.1016/j.ncl.2019.05.002

Zhang, R., Lahens, N.F., Ballance, H.I., Hughes, M.E., Hogenesch, J.B., 2014. A circadian gene expression atlas in mammals: Implications for biology and medicine. Proc. Natl. Acad. Sci. U.S.A. 111, 16219–16224. 10.1073/pnas.1408886111

